# Direct colorimetry of imipenem decomposition as a novel cost effective method for detecting carabapenamase producing bacteria

**DOI:** 10.1101/2022.03.13.484133

**Authors:** Stathis D. Kotsakis, Anastasia Lambropoulou, Georgios Miliotis, Eva Tzelepi, Vivi Miriagou, Leonidas S. Tzouvelekis

**Author notes:** Corresponding author: Dr. Stathis D. Kotsakis, Laboratory of Bacteriology, Hellenic Pasteur Institute.

## Abstract

In the absence of a molecule that would collectively inhibit both metallo-β-lactamases and serine reactive carbapenemases, containment of their genes’ spreading is the main weapon currently available for confronting carbepenem resistance in hospitals. Cost effective methodologies rapidly detecting carbapenemase producing enterobacteria (CPE) would facilitate such measures. Herein a low cost CPE detection method was developed that was based on the direct colorimetry of the yellow shift caused by the accumulation of diketopiperazines – products of the acid catalyzed imipenem oligomerization – induced by carbapenemase action on dense solutions of imipenem/cilastatin. The reactions were studied by spectrophotometry in the visible spectrum using preparations of β-lactamases from the four molecular classes. The effects of various buffers on reactions containing the potent carbapenemases NDM-1 and NMC-A were monitored at 405 nm. Optimal conditions were used for the analysis of cell suspensions and the assay was evaluated using 38 selected enterobacteria including 29 CPE as well as nine carbapenemase-negative strains overexpressing other β-lactamases. The development of the yellow color was specific for carbapenemase containing enzyme preparations and the maximum intensity was achieved in acidic or un-buffered conditions in the presence of zinc. When applied on bacterial cell suspensions the assay could detect CPE with 96.7 % sensitivity and 100 % specificity with results being comparable to those obtained with the CARBA NP technique. Direct colorimetry of carbapenemase-induced imipenem decomposition required minimum reagents while exhibited high accuracy in detecting CPE. Therefore it should be considered for screening purposes after further clinical evaluation.

**Importance:** Currently, spread of multi-drug resistant (MDR) carbapenemase-producing enterobacteria (CPE), mostly in the clinical setting, is among the most pressing public health problems worldwide. In order to effectively control CPE, use of reliable and affordable methods detecting carbapenemase genes or the respective β-lactamases is of vital importance. Herein we developed a novel method, based on a previously undescribed phenomenon, which can detect CPE with few reagents by direct colorimetry of bacterial suspensions and imipenem/cilastatin mixtures.

## Introduction

The development of a single molecule that would collectively inhibit all carbapenemases is a difficult task due to their different reaction mechanisms (zinc dependent; metallo-beta-lactamases or serine reactive; classes A and D). Although molecules with such properties are currently being tested (e.g. cyclic boronates; 1), none has entered the clinical practice and the latest therapeutic β-lactam/β-lactamase inhibitor combinations encompassing the diazabicyclooctanes class of compounds (e.g. avibactam and relebactam) are only active against serine reactive enzymes (2). Given that carbapenemase producing enterobacteria (CPE) are commonly expressing co-resistances to other drugs of choice, the early detection and confinement of their sources is currently the main way that could confront outbreaks of the respective infections in a health-care setting (3).

A high number of CPE diagnostic techniques are currently available (reviewed in references 4, 5). Yet, only a handful are suitable for the screening purposes of an infection control approach. An efficient technique for CPE screening should be high throughput, sensitive and specific, cost effective, able to detect even unknown carbapenemases while also providing information on the reaction mechanism (MβL or serine reactive). The above criteria are simultaneously satisfied by methodologies which detect the carbapenemase activity in dense bacterial suspensions using color development (e.g. RAPIDEC CarbaNP, BlueCarba, beta-CARBA and MAST CARBA PAcE; 6–9). Although the current colorimetric techniques are relatively low cost, the cumulative financial burden during a screening would be still high for a limited-budget setting.

Recently, during the development of a technique that detects the imipenem acidic hydrolysis product using an ion sensitive field effect transistor (10) we have observed that reaction mixtures containing CPE yielded a yellowish color that gradually became more intense - something that did not occur for the carbapenemase negative strains even after prolonged incubation. This phenomenon most likely resulted from the pH drop during imipenem hydrolysis and was due to the complex oligomerization reactions taking place in dense solutions of the compound under acidic conditions yielding chromophoric diketopiperazines (11). Herein we showed that under the experimental conditions of the techniques detecting carbapenemase production utilizing pH changes, a color shift would occur due to imipenem decomposition, even in the absence of an indicator and that this can be used as a cost effective alternative method for CPE screening.

## Materials and Methods

### β-Lactamase preparations

Crude protein extracts containing β-lactamases were prepared from laboratory *E. coli* clones replicating the recombinant plasmids pZE21-*bla*_NDM-1_ (*E. coli* C600Z1), pNTN3-*bla*_NMC-A_ (*E. coli* JM109), pZE21-bla_OXA-48_ (*E. coli* C600Z1), pBC-*bla*_CMY-2_ (*E. coli* DH5α) and pBC-*bla*_CTX-M-15_ (*E. coli* DH5α) overexpressing the NDM-1 (MβL), NMC-A (class A carbapenemase), OXA-48 (class D carbapenemase), CMY-2 (class C β-lactamase) and CTX-M-15 (class A β-lactamase -ESBL) enzymes respectively. In pZE21 clones transcription of the cloned β-lactamase gene was induced by 200 ng/ml anhydrotetracycline while in the remaining plasmids expression was constitutive driven by natural promoters of the genes. Proteins were released through sonication (10) in 50 mM sodium phosphate bufffer pH 7 with the exception of the MβL preparations where a 50 mM HEPES, 50 μΜ ZnSO_4_ pH 7.2 buffer was used. Hydrolysis of imipenem (NDM-1, NMC-A and OXA-48), cephalothin (CMY-2) or cefotaxime (CTX-M-15) was measured by UV spectrophotometry. β-Lactamase concentration in the extracts was estimated using the initial velocities and the published steady state hydrolysis constants (12–16) by the *Michaelis-Menten* equation.

### Bacterial strains and susceptibility testing

A total of 38 non-repetitive enterobacterial strains were used in the study. These included 29 strains producing a carbapenemase and nine strains producing β-lactamases with either marginal or no carbapenem hydrolytic activity (non-carbapenemases). The detailed β-lactamase content of each strain is given in Table 1. Isolates had been previously characterized using phenotypic and molecular techniques (17, 18). The uniqueness of strains belonging in the same species and exhibiting identical β-lactamase content was asserted through restriction fragment length polymorphism analysis using pulsed field gel electrophoresis. All strains had been isolated from clinical settings in Greece, save for the IMP producers that were of environmental origin (19).

**Table 1:** Clinical strains used in the study and absorbance changes at 405 nm during incubation of bacterial suspensions with 5 mg/ml imipenem/cilastatin.

| Strain | β-lactamases | Max $\Delta A_{405\text{ nm}}$ | $t_{\Delta A>0.05}$ (min) | MIC ($\mu\text{g/ml}$) | |
| --- | --- | --- | --- | --- | --- |
|  |  |  |  | Imipenem | Meropenem |
| VIM MβL producers | | | | | |
| *K. pneumoniae* IK14 | VIM-1, CMY-2 type | 0.53±0.01 | 90 | 1 | 0.5 |
| *K. pneumoniae* LA30 | VIM-1, CMY-2 type | 0.57±0.01 | <30 | 8 | 16 |
| *E. coli* TZ116 | VIM-1, CMY-13 | 0.54±0.01 | <30 | 4 | 1 |
| *E. cloacae* SEC6 | VIM-1 | 0.53±0.05 | 120-180 | 1 | 1 |
| *P. mirabilis* EUG91 | VIM-1, VEB-1 | 0.39±0.01 | 120 | 4 | ≤0.125 |
| NDM MβL producers | | | | | |
| *K. pneumoniae* LA26 | NDM-1, CTX-M-15 | 0.42±0.07 | <30 | 16 | 32 |
| *K. pneumoniae* LA27 | NDM-1, CTX-M-15 | 0.48±0.05 | <30 | 8 | 8 |
| *K. pneumoniae* LA28 | NDM-1, CTX-M-15 | 0.63±0.01 | <30 | 8 | 64 |
| *K. pneumoniae* 2489 | NDM-1, CTX-M-15 | 0.41±0.02 | <30 | 32 | 64 |
| *K. pneumoniae* LA | NDM-1, CTX-M-15 | 0.59±0.05 | <30 | 8 | 64 |
| *K. aerogenes* 7362 | NDM-1 | 0.61±0.07 | <30 | | |
| IMP MβL producers | | | | | |
| *K. pneumoniae* 1733 | IMP-4 | 0.53±0.06 | <30 | ND | 1 |
| *E. coli* 1543m1 | IMP-4 | 0.49±0.05 | <30 | ND | 4 |
| *E. coli* 1586m1 | IMP-38 | 0.45±0.05 | <30 | ND | 0.5 |
| *E. cloacae* 1537m | IMP-4 | 0.47±0.03 | <30 | ND | 8 |
| *P. mirabilis* 1539p | IMP-4 | 0.39±0.03 | <30 | ND | 8 |
| KPC-2 class A carbapenemase producers |  |  |  |  |  |
| *K. pneumoniae* E971 | KPC-2, SHV-5 type | 0.49±0.05 | 60 | 1 | 8 |
| *K. pneumoniae* E1370 | KPC-2 | 0.62±0.01 | <30 | 2 | 4 |
| *K. pneumoniae* E1505 | KPC-2 | 0.68±0.04 | <30 | 32 | 16 |
| *E. coli* LAR373 | KPC-2 | 0.71±0.03 | <30 | 2 | 1 |
| *E. coli* LAR152 | KPC-2, CTX-M, OXA-1 | 0.49±0.07 | <30 | 2 | 2 |
| OXA-48 class D carbapenemase producers |  |  |  |  |  |
| *K. pneumoniae* ALX47 | OXA-48, CTX-M-15 | 0.23±0.02 | 180 | 4 | 8 |
| *K. pneumoniae* LAR478 | OXA-48, CTX-M-15 | 0.13±0.01 | 240 | 8 | 8 |
| *K. pneumoniae* AO2085 | OXA-48, CTX-M-15 | 0.31±0.03 | 180 | ND | ND |
| *K. pneumoniae* AO1938 | OXA-48, CTX-M-15 | 0.32±0.03 | 180 | ND | ND |
| *K. pneumoniae* NQ3083 | OXA-48 | 0.57±0.05 | 90 | 8 | 4 |
| *K. pneumoniae* NQ4129 | OXA-48 | 0.47±0.03 | 90 | 4 | 4 |
| *K. pneumoniae* TRK1 | OXA-48, CTX-M-15 | -0.04±0.04 | NA | 8 | 32 |
| *K. pneumoniae* TRK5 | OXA-48, CTX-M-15 | 0.06±0.04 | 360 | 16 | 32 |
| Carbapenemase negative strains |  |  |  |  |  |
| *K. pneumoniae* TZ59 | species specific SHV | -0.002±0.003 | NA | ≤0.125 | ≤0.125 |
| *K. pneumoniae* IPT59 | GES-7, SHV-5 | 0.01±0.03 | NA | 0.25 | ≤0.125 |
| *K. pneumoniae* 17829 | CTX-M-15, SHV-12 | -0.01±0.01 | NA | 16 | 16 |
| *K. pneumoniae* EY-205 | CMY-36, SHV-5 | 0.002±0.03 | NA | 0.25 | ≤0.125 |
| *K. aerogenes* EY-25 | LAT-2+SHV-5 | -0.01±0.03 | NA | 16 | 8 |
| *E. cloacae* WTEY-138 | derepressed AmpC | 0.009±0.1 | NA | 0.5 | ≤0.125 |
| *E. coli* IK33 | CTX-M-15 | 0.003±0.006 | NA | ≤0.125 | ≤0.125 |
| *E. coli* S13 | CTX-M-32 | 0.009±0.005 | NA | ≤0.125 | ≤0.125 |
| *K. aerogenes* EY15 | LAT-2 | 0.007±0.004 | NA | 8 | 4 |

ND: Not determined, NA: Not applicable

Imipenem and meropenem MICs were determined using the microdillution method in Mueller-Hinton broth according to the EUCAST recommendations. Carbapenems were tested at a concentration range from 0.125 to 128 μg/ml.

### Spectrophotometric analyses of imipenem decomposition in the visible

In spectrophotometric analyses a stock solution of 10 mg/ml imipenem – 10 mg/ml cilastatin was used. It was prepared from a generic 500+500 mg imipenem/cilastatin powder for injection containing also 1.6 mmol of sodium bicarbonate (NaHCO_3_). Reconstitution was carried out using either a solution of 0.3 mM zinc sulfate (ZnSO_4_) or de-ionized water and the resulting suspensions were aliquoted and stored at −80 °C until further use.

Acquisition of absorbance spectra was carried out using a HITACH U-2001 UV/Vis double beam spectrophotometer in a quartz cuvette of 1 cm optical path. Each reaction had a volume of 1 ml and was prepared through 1:1 dilution of the imipenem/cilastatin-ZnSO_4_ stock solution in deionized water that resulted in the following composition: 5 mg/ml imipenem, 5 mg/ml cilastatin, 16 mM NaHCO_3_ and 0.15 mM ZnSO_4_ with the pH being 7.2±0.1. Quantities of the β-lactamase preparations were added in the reaction mixture - with the buffering salt included in the crude protein extract having a final concentration of no more than 2 mM - and the spectrum from 342 nm to 1100 nm was scanned at a rate of 800 nm∙min^−1^ at various time intervals. Differential absorption spectra were obtained through subtraction of the initial spectrum from the spectra obtained at each time point. A control reaction lacking a β-lactamase was also performed as above.

The effects of zinc, pH and various buffers on the carbapenemase induced color development were examined using a DYNEX MRX absorbance micro-plate reader. Readings were obtained at 405 nm with the reference filter being set at 630 nm. Here the imipenem/cilastatin-water stock solution was used that was diluted 1:1 either in i) deionized water, ii) 0.1 M 2-(N-morpholino)-ethanesulfonic acid (MES) pH 5.4, iii) 0.1 M 3-(N-morpholino)-propanesulfonic acid (MOPS) pH 6.9, iv) 0.1 M MOPS pH 7.2 or v) 0.1 M Tris/HCl pH 8. The same solutions supplemented with 0.3 mM ZnSO_4_ were also assayed. Reactions were prepared directly on the microplate’s wells and had a volume of 100 μl. The NDM-1 MβL and the NMC-A class A carbapenemases were tested and results were compared with those of control wells.

The effects of NDM-1 and NMC-A carbapenemases on 5 mg/ml of imipenem (imipenem hydrate ≥98%, Cayman Chemicals) solution containing 16 mM NaHCO_3_ and 0.15 mM ZnSO_4_ as well as on 5 mg/ml imipenem – 5 mg/ml cilastatin in 0.15 mM ZnSO_4_ prepared from the brand name Primaxin formulation (Merck, Sharp& Dohme Corp.) were also examined.

### Analysis of bacterial suspensions with the imipenem decomposition method

Dense cell suspensions were prepared by the addition of two full 10 μL plastic inoculation loops (Sarstedt, Germany) of bacteria grown on Tryptone Soya Agar (TSA; OXOID-Thermo Scientific, UK), supplemented with 0.3 mM ZnSO_4_, into 400 μL of H_2_O. For each strain, 50 μL of this suspension were added into four wells of a 96-well micro-plate (polysterene flat bottom clear wells; Greiner, Germany). Fifty microliters of the imipenem/cilastatin-ZnSO_4_ stock solution were added in two of the above wells while in the remaining two, introduced for absorbance correction, the same volume of a 0.3 mM ZnSO_4_ solution was added (control 1). Wells containing the Imipenem/cilastatin-ZnSO_4_ (control 2) and ZnSO_4_ solutions (control 3) diluted 1:1 with H_2_O were also included as controls. The plates were incubated at 37 °C and the absorbance was measured using a DYNEX MRX micro-plate reader at various time points. The absorbance of the wells containing the mixtures of bacterial suspensions with imipenem/cilastatin were corrected by subtracting that of control 1 and control 2 (control 3 corrected) wells. Each experiment was performed in triplicate. Estimation of a threshold of absorbance increase in order to characterize a strain as a carbapenemase producer was carried out through Receiver Operating Characteristic (ROC) analysis with Prism v. 8.0. Absorbance changes documented in experiments of carbapenemase negative strains were grouped in the “Control column” and those of positive strains in the “Patients” column. An absorbance increase greater than 0.0456 yielded 96.55% sensitivity (95% Confidence Interval, CI: 82.82 – 99.82%) and 100% specificity (95% CI: 97.23 – 100.00%). Hence the threshold was set at 0.05 of absorbance increase.

The effect of metal chelation on color development was assessed by the addition of EDTA (0.5 M pH 8) in the bottom of the wells before the various reaction components and by preparing the bacterial suspensions in EDTA containing solutions. The concentrations of EDTA included in the reaction during preliminary experiments were 10 and 15 mM with the former being selected as optimum. In these experiments bacteria grown on TSA without zinc supplementation were also tested.

### Comparisons with the CARBA NP technique

Direct colorimetry was compared with the commercial pH indicator colorimetric technique RAPIDEC CARBANP (bioMerieux, France). For these comparisons we included strains expected to cause sensitivity and specificity issues in CPE detection techniques. Bacteria were grown on TSA containing 0.3 mM ZnSO_4_ at 37 °C for 16 h and the assay was performed and interpreted according to the manufacturer’s instructions.

## Results and discussion

### Spectrophotometric analyses of β-lactamase – imipenem/cilastatin mixtures in the visible

β-Lactam hydrolysis is accompanied by shifts in absorption in the UV spectrum due to the opening of the four member ring. The fact that carbapenemase action on imipenem solutions in the absence of a pH buffer leads to absorbance changes in the visible region of the light spectrum, as the yellow color development indicated, prompted us to study the phenomenon through spectrophotometry.

The differential absorption spectra (Figure 1A) in carbapenemase containing reactions showed the accumulation of species that absorb in the violet region. The efficient carbapenemases NDM-1 and NMC-A assayed at nanomolar quantities induced shifts which were apparent after 15 minutes. The less potent OXA-48 required longer reaction times and sub-micromolar quantities in order to observe the absorbance increases in the violet region (Figure 1A, upper panel). On the other hand, CMY-2 and CTX-M-15 – enzymes that do not exhibit meaningful imipenemase activity – yielded only minor absorbance increases after three hours, similarly with what was observed in the control reaction containing solely imipenem/cilastation (Figure 1A, lower panel). Therefore, the color shift was a phenomenon that was specifically observed for carbapenemases.

**Figure 1:**
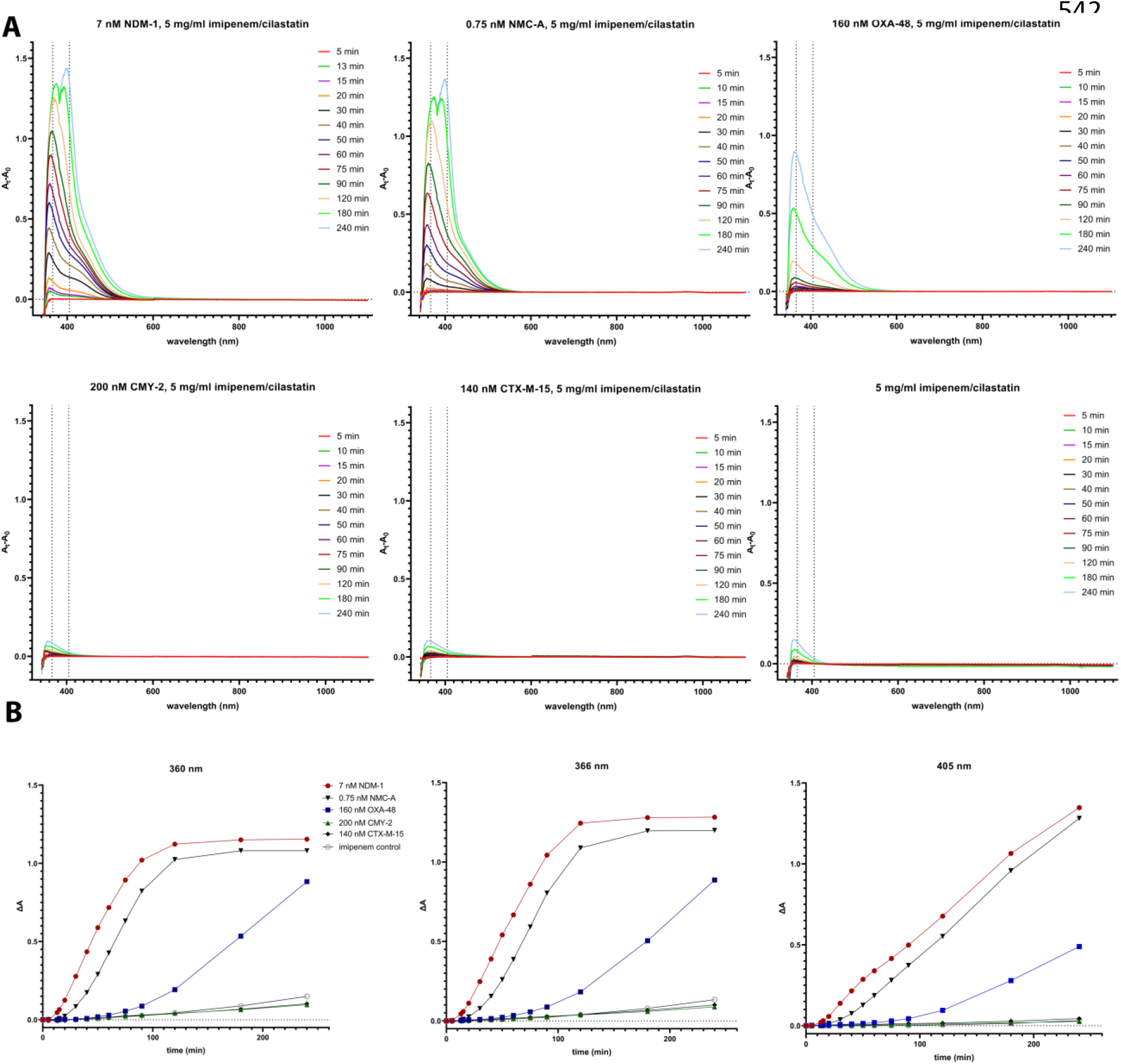
**A)** Differential absorption spectra in the visible spectrum during incubation of various β-lactamases with 5 mg/ml imipenem/cilastatin. The efficient carbapenemases NDM-1 and NMC-A induced sharp absorbance shifts followed by the less potent OXA-48. Enzymes not exhibiting imipinemase activity caused marginal absorbance changes that were comparable to that of the control reaction. **B)** Absorbance increases in three wavelengths corresponding to the detected peaks in differential absorption spectra. Monitoring the absorbance increase at 405 nm can clearly differentiate carbapenemases from non-carbapenemases.

Multiple peaks were apparent with the λ_max_ initially being 360 nm and then increased with time up to 370 nm. In the highly efficient NDM-1 and NMC-A, after three hours of incubation a second peak became predominant in the area of 400 nm while absorbance in the previous peak remained stable (Figure 1A). The above data indicated that the reactions taking place during the action of carbapenemases resulted in the formation of more than one chromogenic products. By plotting the absorbance in various wavelengths it was apparent that the carbapenemase activity could also be detected at 405 nm, though requiring longer reaction times (Figure 1B). Hence, the carbapenemase induced color shift could be quantified in a Clinical Microbiology laboratory through the widely available micro-plate absorbance readers instead of a UV/Vis spectrophotometer used here.

### Effects of various solutions on carbapenemase induced color formation

The effects of different buffers at various pH values in the presence and absence of zinc(II) on the occurrence of the yellowish color were assessed. In NDM-1 containing reactions the color development was dependent on zinc, especially at the acidic pH of the MES buffer as well as in un-buffered conditions, contrary to NMC-A and control experiments (Figure 2A). Coloration induced by NMC-A was dependent on the pH and the buffering capacity of the solution. The highest absorbance increases were observed in reactions which did not contain a buffering salt (i.e. Η_2_Ο or 15 mM ZnSO_4_ reactions) and in the MES buffer at pH 5.4 (Figure 2A; right graph column). In the presence of MOPS, phosphate and Tris buffers the color development was significantly attenuated as the alkalinity of the reaction environment increased (Figures 2A and 2B). Similar observations were made for NDM-1 in zinc supplemented solutions (Figures 2A and 2B). In control reactions containing only imipenem a moderate absorbance increase was observed in the acidic MES buffer irrespective of zinc ions with the remaining solutions being inert (Figure 2A; left graph column). The specific requirement for Zn(II) in MβL reactions provided additional evidence that the phenomenon is indeed induced by the enzymatic hydrolysis of imipenem. It has been shown that low pH has a detrimental effect on MβL activity - probably due to the protonation of Asp120 of the second Zn(II) binding site that results in loss of one of the zinc ions – and that this can be countered by zinc supplementation of the reaction buffer (20, 21).

**Figure 2:**
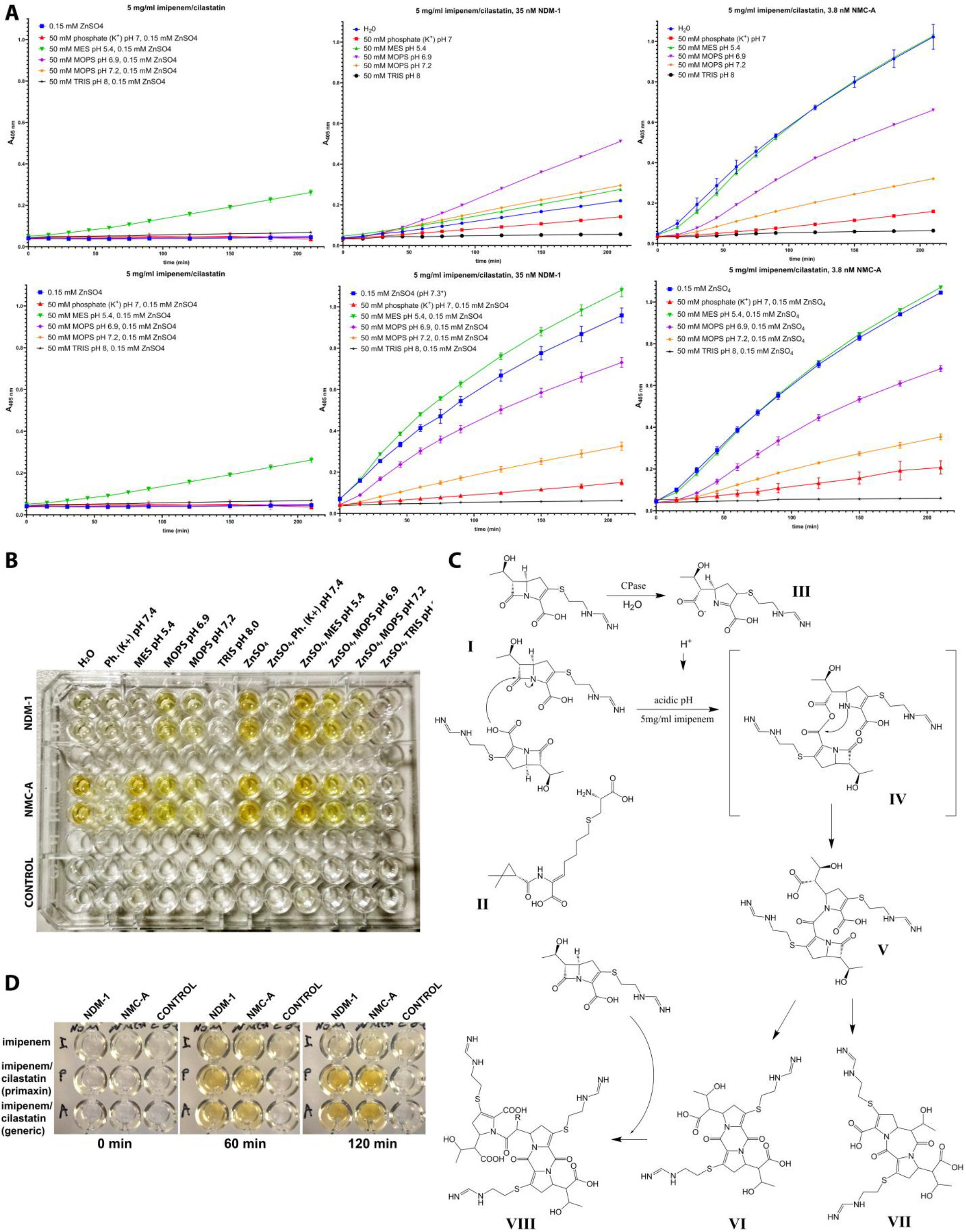
**A)** Effects of various buffers on the color development induced by NDM-1 and NMC-A in the absence (upper panel) and presence (lower panel) of zinc cations estimated by absorbance measurements at 405 nm. In NDM-1 containing reactions (medium column) color development is significantly enhanced by zinc sulfate at a concentration of 0.15 mM while the class A carbapenemase NMC-A (right column) yielded equivalent signals in both conditions. Color development was extended in acidic pH and in un-buffered conditions, with high alkalinity attenuating the reaction. In control reactions lacking a carbapenemase (left column) an absorbance increase was observed in the presence of 50 mM MES buffer pH 5.4. **B)** Color development in the above conditions as documented at the final point (t= 240 min). **C)** A likely molecular mechanism for the carbapenemase induced yellow color. **CPase:** carbapenemase, **I:** imipenem, **II:** cilastatin, **III:** hydrolyzed imipenem. Compounds **VI** and **VIII**, containing a diketopiperazine ring, exhibit a λ_max_ at 360 nm and form yellow colored solutions (11). **D)** Effect of NDM-1 and NMC-A on pure imipenem and on the brand name imipenem/cilastatin formulation (Primaxin) in comparison with the generic imipenem/cilastatin formulation used in the study under the same conditions. Imipenem/cilastatin reactions yielded stronger and more stable yellow color compared to pure imipenem.

The dependence of the carbapenemase induced color development on acidic pH or the absence of a buffering agent provided some evidence on the likely molecular bases of the phenomenon. It is known from stability studies of imipenem and the imipenem/cilastatin formulation that the compound decomposes in acidic pH at concentrations ≥1 mg/ml through complex oligomerization reactions that lead to the formation of diketopiperazines yielding yellow colored solutions (11, 22–25). Hence, a possible explanation for our data would be that as the enzymes hydrolyze the β-lactam ring of imipenem and the acidic hydrolysis product is accumulated, the pH is decreased triggering thus secondary decomposition reactions leading to the formation of chromogenic diketopiperazines (Figure 2C - compounds VI and VIII; 11).

The developed assay requires increased quantities of the substrate and hence, to decrease the cost, we have used a commercially available generic imipenem/cilastatin formulation. In order to assert that the observed phenomenon is governed by the above mechanism we assayed pure imipenem and the brand-name imipenem/cilastatin formulation (Primaxin). After 60 minutes of incubation the yellow color was developed in all reactions, with those of imipenem/cilastatin yielding stronger signals (Figure 2D). At 2 hours though, the color in the imipenem solution started to fade, indicating consumption of the chromophore product, in contrast to the imipenem/cilastatin solutions (Figure 2D). The above results suggested that imipenem oligomerization caused by the acidification induced by the action of carbapenemases may indeed be the reason for the color development, at least in the initial reactions, with cilastatin having a yet unknown key role.

### Developement of a CPE screening tool

As assays with enzyme preparations indicated that the color development due to imipenem decomposition was specific for carbapenemases we subsequently explored the use of this method as a diagnostic tool by testing cell suspensions of clinical isolates. Although the color shift was visually detectable we quantified it through absorbance measurements at 405 nm using a micro-plate reader to improve objectivity.

The majority of the MβL producing enterobacteria exhibited rapid color shifts that were also reflected on the measured absorbance (Figure 3; Table 1). The weakest responses were observed with VIM-1 expressing strains with three of them requiring more than 60 minutes incubation in order the yellow color to develop (Figure 3). Nonetheless, all 15 ΜβL producers yielded high intensity end-point coloration with the maximum absorbance at 405 nm being in the range of 0.39 to 0.43 units (Figure 3; Table 1). Fast color development was also evident for all KPC-2 class A carbapenemase producers tested with the maximum absorbance values ranging from 0.49 to 0.71 (Figure 3; Table 1). Production of the less efficient OXA-48 class D carbapenemase required longer incubation times in order to be detected through the imipenem decomposition method with the yellow color developing after 90 to 120 minutes (Figure 3). Furthermore two strains, isolated in the initial stages of the OXA-48 epidemic in the Near East, yielded marginal or no color shifts (*K. pneumoniae* TRK-5 and TRK-1; Figure 3). These strains were found negative with CARBA NP (Table 2). In the six OXA-48 producing strains that yielded a response the maximum absorbance varied between 0.06 and 0.57. The nine isolates not producing a carbapenemase but overexpressing other β-lactamases did not yield any coloration even after six hours of incubation (Figure 3). The maximum absorbance values observed for these strains ranged between −0.002 to 0.01 units (Table 1). By applying the threshold estimated trough ROC analysis the method could detect 16 out of 16 of the MβL strains within <30-180 minutes, 5/5 of KPC-2 producers in less than 30 minutes, and 6/7 of OXA-48 isolates in 90 to 360 minutes while it excluded all the non carbapenemase producers as negatives (Table 1). Of note, two of the carbapenemase negative isolates (*K. pneumoniae* EY-205 and 17829) gave false positive results when analyzed with the CARBA NP technique (Table 2). The obtained data indicated that direct colorimetry could detect CPE with 96.4% sensitivity (1/28 false negatives) and 100% specificity (0/9 false positives).

**Table 2:**
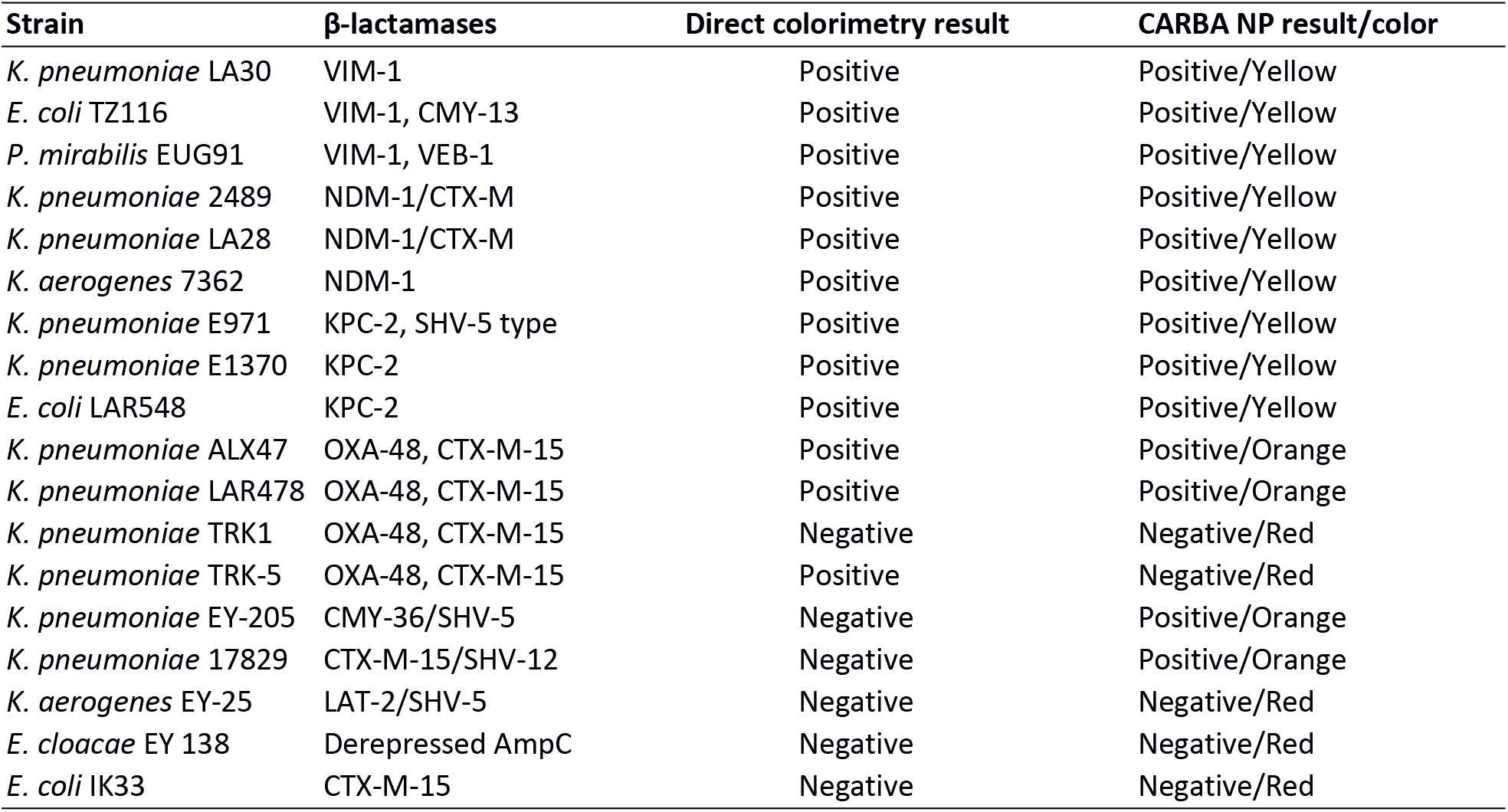
Comparison of the direct colorimetry method with RAPIDEC CARBA NP for selected strains

**Figure 3:**
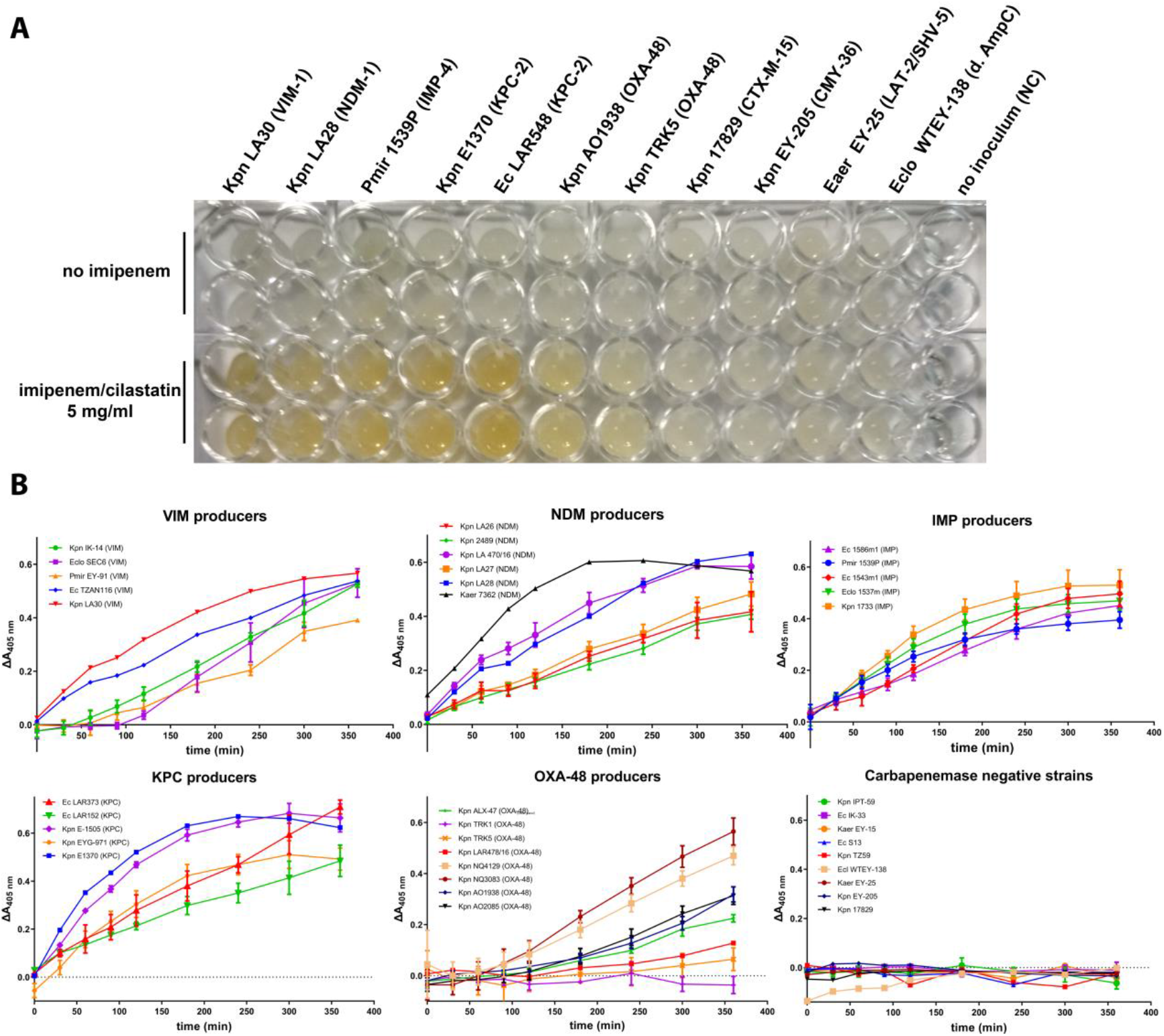
Application of the imipenem decomposition method on bacterial suspensions. **A)** Color development at the end point (t = 360 min) of enterobacterial strains’ cell suspensions producing various types of carbapenemases and β-lactamases with no imipenemase activity. **B)** Absorbance changes at 405 nm during the course of six hours in imipenem/cilastatin-bacterial suspensions mixtures of the strains assayed in the study. MβL and KPC producing strains yielded strong signals that could be detected as early as 30 minutes. *K. pneumoniae* producing the OXA-48 class D carbapenemase yielded weaker responses with one strain being identified as negative. The nine non-carbapenemase producers did not cause any color shift even after six hours of incubation.

The ability of this method to discriminate between ΜβL and serine reactive carbapenemase producers was assessed using EDTA as a chelating agent. Preliminary experiments were performed using 10 and 15 mM EDTA in the reaction mixtures. At these concentrations the color formation observed in MβL producing strains was attenuated but a quenching in the coloration induced by KPC-2 producers was observed, probably due to an increase of the solution’s alkalinity (Supplemental Figure 1). Hence, 10 mM was selected as an optimal EDTA concentration. EDTA could efficiently inhibit the yellow color induced by suspensions of NDM producing bacteria as well as of some VIM and IMP producing strains with insignificant effects on signals obtained from serine reactive carbapenemase producers (Figure 4). Color formation in the presence of EDTA was evident with a VIM-1 producing *K. pneumoniae* (Kpn LA30) and a *P. mirabilis* strain expressing the IMP-4 enzyme (Figure 4A, right panel). Considering that the yellow color is formed indirectly through secondary reactions and not due to the direct action of carbapenemases, then any residual imipenem hydrolysis may initiate the cascade leading to a positive result. In order to overcome this we performed the same experiments with the bacterial suspensions being prepared in EDTA which was then mixed with the imipenem/cilastatin solution. This modification permitted the inhibition of *K. pneumoniae* LA30 reactions but not those of the *P. mirabilis* IMP-4 strain that was still able to yield a strong coloration (Figure 4A and C). The above may be due to the relative resistance of IMP enzymes to the action of EDTA combined with increased levels of the enzyme in the bacterium’s periplasm.

**Figure 4:**
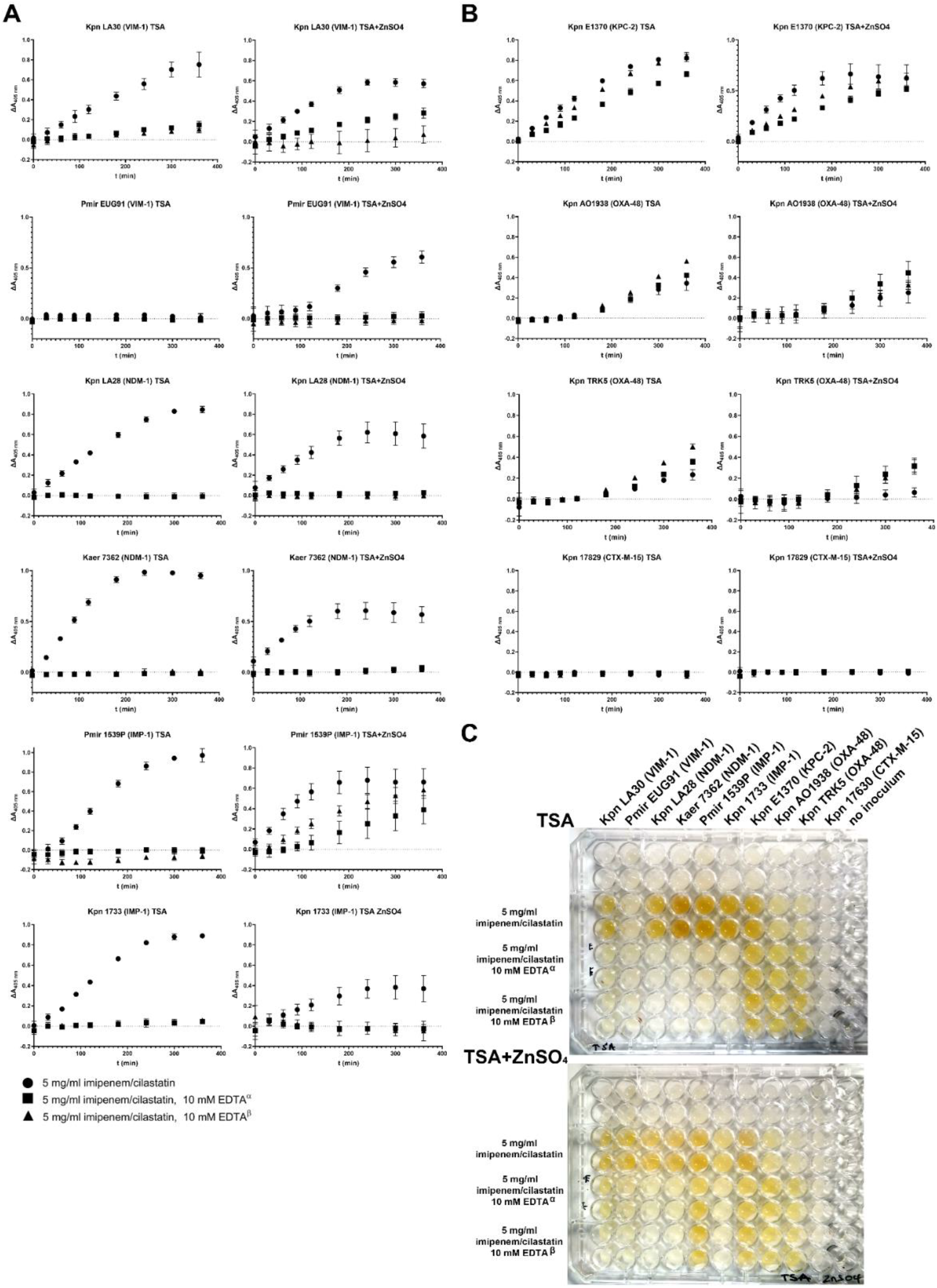
Differentiation of MβL and class A carbapenemase producers through the use of EDTA and effects of zinc cation supplementation of the growth medium. **(A)** Time courses of absorbance changes at 405 nm of MβL producing strains in the presence and absence of 10 mM EDTA when bacteria were cultured without (left panel) and with zinc supplementation (right panel) in the TSA medium. Experiments were performed with EDTA added in the wells prior to reactants’ addition (black squares, α) or with bacterial cells suspended in a 20 mM EDTA solution (black triangles, β). EDTA inhibited the decomposition of imipenem in the majority of MβL strains when they were cultivated in the absence of zinc. *P. mirabilis* EUG91 did not yield any signal when grown on plain TSA. **(B)** Effects of EDTA and growth on zinc supplemented TSA on reactions containing strains producing serine reactive β-lactamases. **(C)** Coloration observed in the above experiments after six hours of incubation.

Indeed, as the strains were grown on zinc supplemented media, in order to assert high periplasmic levels of fully functional ΜβLs, the observed resistance of the color formation to EDTA inhibition may be due to increased enzyme quantities released in the solution. In order to assess this we performed the same set of experiments with strains grown on plain TSA. The results showed that without zinc supplementation, the color development could be inhibited by EDTA in the majority of the MβL producing strains (Figure 4A, left panel). However *P. mirabilis* EUG91, that produces low quantities of VIM-1 (10), failed to give a positive reaction even after six hours of incubation. Thus, EDTA inhibition could be used for the identification of MβL producers with the above method when bacteria are cultured without excess zinc but this would reduce the sensitivity for some strains exhibiting low levels of functional periplasmic MβLs.

The color development induced by suspensions of serine reactive carbepenemase producing enterobacteria remained relatively unaffected by EDTA (Figure 4B and C). It should be noted that in OXA-48 producers the chelator increased the color intensity, probably through the release of more enzyme in the solution facilitated by its detrimental effect on cell wall integrity (Figure 4B). Zinc supplementation of the growth media seemed to affect the chromogenic reaction of positive strains yielding systematically lower end point absorbance increases even for the MβL producing isolates (*Δ*OD^TSA-max, TSA+ZnSO4-max^ = 0.26±0.12, paired t-test p<0.001). Zinc ions are known to have a variety of effects on bacterial physiology and therefore we cannot yet provide an explanation for the above observation. Strains not producing a carbapenemase remained negative under all the employed modifications of the method.

Based on these data the use of direct colorimetry combined with various chelating agents in order to distinguish MβL from serine carbapenemase producers warrants further study.

## Conclusions

Imipenem is the first clinically used carbapenem that exhibits increased stability to aminolysis compared to its natural counterpart thienamycin. Yet, the molecule still possesses an inherent instability in aqueous solutions as compared to newer carbapenems (e.g. meropenem; 27, 28). Indeed its decomposition in low pH results in complex degradation products not observed in the other members of the group (29). Herein we showed that the accumulation of chromophoric decomposition products of imipenem in the acidic conditions induced by the action of carbapenemases can be used for the specific detection of CPE directly from bacterial suspensions.

Direct colorimetry required minimum reagents i.e. a solution of 10 mg/ml imipenem-10 mg/ml cilastatin containing 0.3 mM zinc sulfate and can be prepared from any imipenem/cilastatin powder formulation for injection used in hospitals. The development of yellow color can be followed either through manual inspection or with a microplate reader capable of measuring absorbance at 405 nm that would increase both through-put and objectivity. The cost of the method would be significantly lower (>100 fold) compared to the current commercial CPE detection colorimetric techniques, considering that a 500 mg imipenem/500 mg cilastatin vial (valued at 6.00 €, Greek market consumer prices; 30) would be sufficient for 500 reactions. Moreover, as direct colorimetry exhibited high accuracy in detecting CPE and discriminating non-carbapenemase producers, it fulfills the requirements of a successful CPE screening technique and merits further evaluation in a variety of clinical settings.

## Aknowledgements

We thank Dr. Angeliki Mavroidi, Department of Microbiology, Konstantopouleio-Patission General Hospital of Athens Greece, for providing *K. pneumoniae* OXA-48 producing strains as well as Dr. Monika Dolejska and Dr. Iva Kutilova, Department of Biology and Wildlife Diseases, University of Veterinary and Pharmaceutical Sciences Brno Czech Republic, for providing the IMP producing enterobacterial strains.

## References

1. Cahill ST, Cain R, Wang DY, Lohans CT, Wareham DW, Oswin HP, Mohammed J, Spencer J, Fishwick CW, McDonough MA, Schofield CJ, Brem J. 2017. Cyclic boronates inhibit all classes of beta-lactamases. Antimicrob Agents Chemother 61.

2. Bush K, Bradford PA. 2019. Interplay between beta-lactamases and new beta-lactamase inhibitors. Nat Rev Microbiol 17:295–306.

3. Tzouvelekis LS, Markogiannakis A, Psichogiou M, Tassios PT, Daikos GL. 2012. Carbapenemases in Klebsiella pneumoniae and other Enterobacteriaceae: an evolving crisis of global dimensions. Clin Microbiol Rev 25:682–707.

4. Aguirre-Quinonero A, Martinez-Martinez L. 2017. Non-molecular detection of carbapenemases in Enterobacteriaceae clinical isolates. J Infect Chemother 23:1–11.

5. Bilozor A, Balode A, Chakhunashvili G, Chumachenko T, Egorova S, Ivanova M, Kaftyreva L, Koljalg S, Koressaar T, Lysenko O, Miciuleviciene J, Mandar R, Lis DO, Wesolowska MP, Ratnik K, Remm M, Rudzko J, Roop T, Saule M, Sepp E, Shyshporonok J, Titov L, Tsereteli D, Naaber P. 2019. Application of molecular methods for carbapenemase detection. Front Microbiol 10:1755.

6. Nordmann P, Poirel L, Dortet L. 2012. Rapid detection of carbapenemase-producing Enterobacteriaceae. Emerg Infect Dis 18:1503–7.

7. Pires J, Novais A, Peixe L. 2013. Blue-carba, an easy biochemical test for detection of diverse carbapenemase producers directly from bacterial cultures. J Clin Microbiol 51:4281–3.

8. Bernabeu S, Dortet L, Naas T. 2017. Evaluation of the beta-CARBA test, a colorimetric test for the rapid detection of carbapenemase activity in Gram-negative bacilli. J Antimicrob Chemother 72:1646–1658.

9. Rezzoug I, Emeraud C, Sauvadet A, Cotellon G, Naas T, Dortet L. 2021. Evaluation of a colorimetric test for the rapid detection of carbapenemase activity in Gram negative bacilli: the MAST(R) PAcE test. Antimicrob Agents Chemother doi:10.1128/AAC.02351-20.

10. Kotsakis SD, Miliotis G, Tzelepi E, Tzouvelekis LS, Miriagou V. 2021. Detection of carbapenemase producing enterobacteria using an ion sensitive field effect transistor sensor. Sci Rep 11:12061.

11. Ratcliffe RW, Wildonger KJ, Di Michele L, Douglas AW, Hajdu R, Goegelman RT, Springer JP, Hirshfield J. 1989. Studies on the structures of imipenem, dehydropeptidase I-hydrolyzed imipenem, and related analogs. The Journal of Organic Chemistry 54:653–660.

12. Docquier JD, Calderone V, De Luca F, Benvenuti M, Giuliani F, Bellucci L, Tafi A, Nordmann P, Botta M, Rossolini GM, Mangani S. 2009. Crystal structure of the OXA-48 beta-lactamase reveals mechanistic diversity among class D carbapenemases. Chem Biol 16:540–7.

13. Hachler H, Kotsakis SD, Tzouvelekis LS, Geser N, Lehner A, Miriagou V, Stephan R. 2013. Characterisation of CTX-M-117, a Pro174Gln variant of CTX-M-15 extended-spectrum beta-lactamase, from a bovine Escherichia coli isolate. Int J Antimicrob Agents 41:94–5.

14. Kotsakis SD, Papagiannitsis CC, Tzelepi E, Tzouvelekis LS, Miriagou V. 2009. Extended-spectrum properties of CMY-30, a Val211Gly mutant of CMY-2 cephalosporinase. Antimicrob Agents Chemother 53:3520–3.

15. Marcoccia F, Bottoni C, Sabatini A, Colapietro M, Mercuri PS, Galleni M, Kerff F, Matagne A, Celenza G, Amicosante G, Perilli M. 2016. Kinetic Study of Laboratory Mutants of NDM-1 Metallo-beta-Lactamase and the Importance of an Isoleucine at Position 35. Antimicrob Agents Chemother 60:2366–72.

16. Mariotte-Boyer S, Nicolas-Chanoine MH, Labia R. 1996. A kinetic study of NMC-A beta-lactamase, an Ambler class A carbapenemase also hydrolyzing cephamycins. FEMS Microbiol Lett 143:29–33.

17. Kotsakis SD, Petinaki E, Scopes E, Siatravani E, Miriagou V, Tzelepi E. 2013. Laboratory evaluation of Brilliance CRE Agar for screening carbapenem-resistant Enterobacteriaceae: Performance on a collection of characterised clinical isolates from Greece. J Glob Antimicrob Resist 1:85–90.

18. Voulgari E, Miliotis G, Siatravani E, Tzouvelekis LS, Tzelepi E, Miriagou V. 2020. Evaluation of the performance of Acuitas(R) Resistome Test and the Acuitas Lighthouse(R) software for the detection of beta-lactamase-producing microorganisms. J Glob Antimicrob Resist 22:184–189.

19. Dolejska M, Masarikova M, Dobiasova H, Jamborova I, Karpiskova R, Havlicek M, Carlile N, Priddel D, Cizek A, Literak I. 2016. High prevalence of Salmonella and IMP-4-producing Enterobacteriaceae in the silver gull on Five Islands, Australia. J Antimicrob Chemother 71:63–70.

20. Davies AM, Rasia RM, Vila AJ, Sutton BJ, Fabiane SM. 2005. Effect of pH on the active site of an Arg121Cys mutant of the metallo-beta-lactamase from Bacillus cereus: implications for the enzyme mechanism. Biochemistry 44:4841–9.

21. Rasia RM, Vila AJ. 2002. Exploring the role and the binding affinity of a second zinc equivalent in B. cereus metallo-beta-lactamase. Biochemistry 41:1853–60.

22. Smith GB, Dezeny GC, Douglas AW. 1990. Stability and kinetics of degradation of imipenem in aqueous solution. J Pharm Sci 79:732–40.

23. Smith GB, Schoenewaldt EF. 1981. Stability of N-formimidoylthienamycin in aqueous solution. J Pharm Sci 70:272–6.

24. Testa B, Mayer JM. 2003. Hydrolysis in drug and prodrug metabolism: chemistry, biochemistry, and enzymology. Wiley-VCH, Zürich; Weinheim.

25. Zaccardelli DS, Krcmarik CS, Wolk R, Khalidi N. 1990. Stability of imipenem and cilastatin sodium in total parenteral nutrient solution. JPEN J Parenter Enteral Nutr 14:306–9.

26. Blencowe DK, Morby AP. 2003. Zn(II) metabolism in prokaryotes. FEMS Microbiol Rev 27:291–311.

27. Takasu Y, Yoshida M, Tange M, Asahara K, Uchida T. 2015. Prediction of the stability of meropenem in intravenous mixtures. Chem Pharm Bull (Tokyo) 63:248–54.

28. Testa B, Mayer JM. 2003. Hydrolysis in drug and prodrug metabolism: chemistry, biochemistry, and enzymology. Wiley-VCH, Zürich; Weinheim.

29. Takeuchi Y, Sunagawa M, Isobe Y, Hamazume Y, Noguchi T. 1995. Stability of a 1 beta-methylcarbapenem antibiotic, meropenem (SM-7338) in aqueous solution. Chem Pharm Bull (Tokyo) 43:689–92.

30. Galinos. 2022. Greek Pharmaceuticals Guide, Imipenem+Cilastatin. https://www.galinos.gr/web/drugs/main/drugs/imipenem-cilastatin. Accessed 12 March.

