## Supplemental Figure 1 for "Direct colorimetry of imipenem decomposition as a novel cost effective method for detecting carabapenamase producing bacteria"

**Supplemental Figure 1:** Preliminary experiments of the EDTA effect on imipenem decomposition induced by enterobacterial strains. Strains were grown on Tryptone Soy Agar supplemented with 0.3 mM $ZnSO_4$. A quantity of 0.5 M EDTA solution pH 8 was added in the bottom of the wells before the addition of reactants in order to give final concentrations of 10 and 15 mM at 100 µL. Absorbance was measured as described in the Materials and Methods section of the manuscript. **(A)** Absorbance changes during the time course. **(B)** Coloration observed at end point (300 min).

**A**

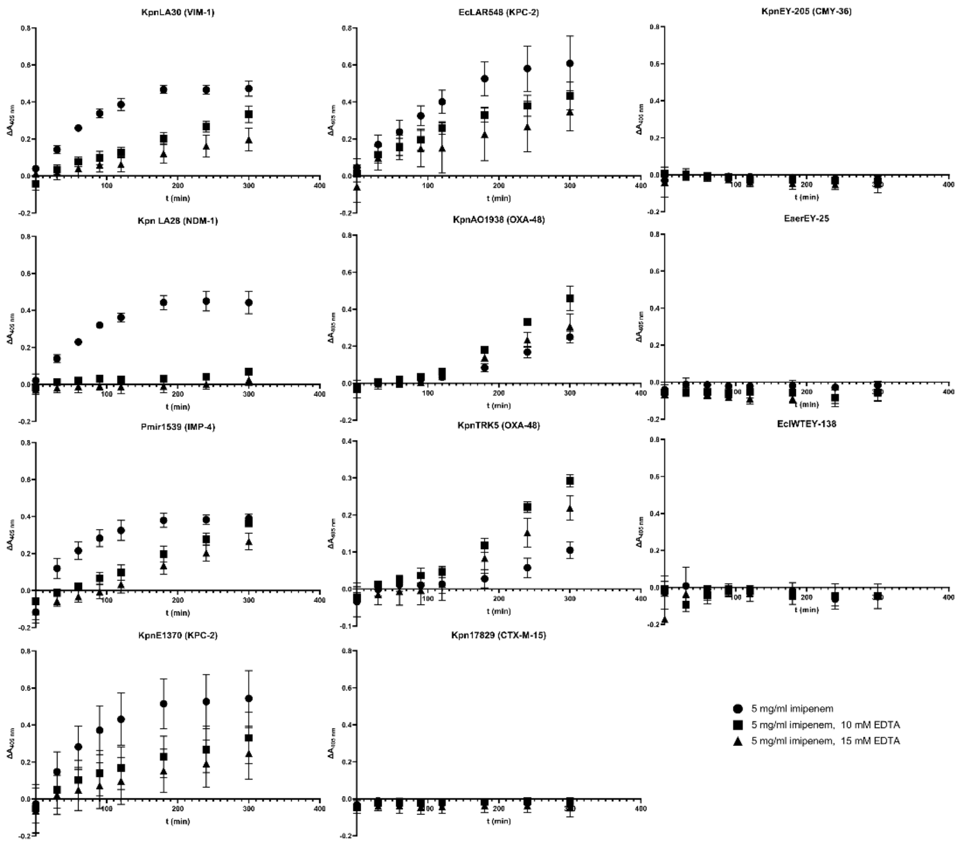


**B**

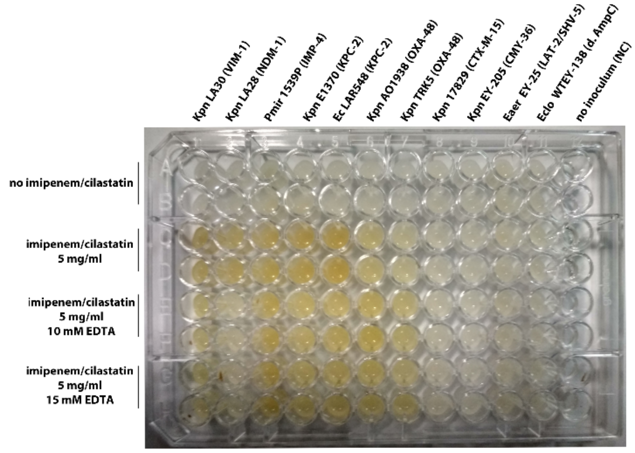
